# Molecular analyses of individual variation in honey bee sociability

**DOI:** 10.1101/2024.12.22.629994

**Authors:** Ian M. Traniello, Arian Avalos, Michael J. M. Gachomba, Tim Gernat, Zhenqing Chen, Amy C. Cash-Ahmed, Adam R. Hamilton, Jennifer L. Cook, Gene E. Robinson

**Author notes:** Corresponding authors. IMT; GER. Lewis-Sigler Institute for Integrative Genomics, Princeton University, Princeton, NJ.

## Abstract

Individual variation in sociability is a central feature of every society. This includes honey bees, with some individuals well-connected and sociable, and others at the periphery of their colony’s social network. However, the genetic and molecular bases of sociability are poorly understood. Trophallaxis – a behavior involving sharing liquid with nutritional and signaling properties - comprises a social interaction and a proxy for sociability in honey bee colonies: more sociable bees engage in more trophallaxis. Here we identify genetic and molecular mechanisms of trophallaxis-based sociability by combining genome sequencing, brain transcriptomics, and automated behavioral tracking. A genome-wide association study (GWAS) identified 18 single-nucleotide polymorphisms (SNPs) associated with variation in sociability. Several SNPs were localized to genes previously associated with sociability in other species, including in the context of human autism, suggesting shared molecular mechanisms of sociability. Variation in sociability also was linked to differential brain gene expression, particularly genes associated with neural signaling and development. Using comparative genomic and transcriptomic approaches, we also detected evidence for divergent mechanisms underpinning sociability across species, including those related to reward sensitivity and encounter probability. These results highlight both potential evolutionary conservation of the molecular roots of sociability and points of divergence.

## Introduction

Consistent variation among individuals in the tendency to engage one another is a central feature of all animal societies. “Sociability,” which broadly characterizes associative interactions outside of the context of aggression or courtship [1], falls on a spectrum, with hypo- and hyper-sociable individuals flanking a “normal” intermediate range [2]. There are many causes of this variation, stemming from a combination of labile drivers like motivational state [3], social status [4], and previous experience [5,6], as well as heritable factors that drive lifelong positive or negative inclinations toward social interactions [7].

As a trait, sociability arises from the integration of socially responsive neural circuitry, physiology, and experience, all of which contribute to how an animal interprets and responds to social cues [8–10]. Detailed study of vertebrate brain evolution has outlined a “social behavior network” that regulates multiple forms of social behavior [11], acting in concert with the mesolimbic reward system to influence how an animal makes decisions in the context of various social situations [2,12,13].

Social insects, including the western honey bee (*Apis mellifera*), have tiny and very differently organized nervous systems relative to vertebrates, but they also form large and intricate societies whose ecological and evolutionary success are tightly linked to individual- and group-level sociability [14,15]. Honey bees form large colonies of tens of thousands of individuals and are a powerful model for further understanding the relationship between genes and sociability. The bulk of the colony is composed of sterile workers for whom task performance is influenced by a combination of individual age, genetics, and the needs of the colony [16–18]. Honey bee workers lack morphologically distinct castes, and each individual maintains the ability to perform all tasks in the colony other than reproduction required for colony growth and development. The molecular mechanisms underlying specific behavioral states like brood care (“nursing”), guarding the hive entrance, or foraging outside for floral resources are sensitive to environmental demands, with clear individual differences in the likelihood of performing certain tasks as well as predisposition toward social contact [5]. We used honey bees to study genetic and molecular components of sociability, including comparative genomic analyses.

Comparative genomics offers a toolkit for exploring the extent to which analogous behaviors, like the tendency to engage socially with others, have similar molecular representations in the brain across disparate species [19,20]. While a behavioral trait may be similarly expressed across species, whether that trait also has a common molecular basis requires observation of similarities across deeper layers of biological organization, including at the level of the genome and/or transcriptome. This approach allows cognitive scientists to better understand the development of the “social brain,” bypassing the need to compare distinct brain anatomies, which is often experimentally challenging, in favor of studying their more quantifiable molecular building blocks. In addition, brain-wide spatial localization of homologous genes or proteins supporting analogous behaviors across species can clarify how vastly different neuroanatomies can support analogous forms of social life [13,21,22]. Drawing from studies in social insects, variation in gene sequence and expression profile have suggested some degree of conservation between vertebrates and invertebrates in the neurogenomic architecture of social behavior, despite ∼600 million years divergence from humans [20,23,24].

Sociability is thought to have a heritable component. In humans, for example, structural variation in the 7q11.23 genomic locus can dramatically affect the developmental trajectory of the social brain, causing either lifelong hypo- or hyper-sociability [7]. Such structural variants affect many aspects of an individual’s neurobiology and physiology, making it challenging to pinpoint direct genetic effects on the propensity to engage in social interactions within a typical range of individual variation. Nevertheless, such findings indicate the existence of heritable molecular substrates that mediate the intensity of sociability.

One limitation in the study of vertebrate sociability is related to difficulties in tracking many individuals over long periods of time – a major challenge in relating genotype and social phenotype in humans [25]. It is similarly difficult to disentangle sociability from related traits, like locomotion, that contribute to overall activity rate: an individual may interact with others more because they are motivated to engage such interactions, because they are simply more physically active and therefore more likely to engage others by chance, or both. But, in some species, it is not possible to characterize individuals across multiple social phenotypic vectors. This issue has been largely overcome for social insects, as recent advances in automated tracking now allow for behavioral monitoring of each individual in a colony over long portions of their lives [26–29]. These advances permit finely resolved comparisons of social behavior, brain, and genome in a naturalistic setting.

In addition to task-related behaviors, honey bees also frequently engage in mouth-to-mouth sharing of liquid containing nutritive and signaling components in a stereotyped process called trophallaxis. Automated behavioral tracking of barcoded bees has shown consistent variation in trophallaxis frequency to be a robust proxy for several aspects of an individual’s life history, from task specialization to disease status [5,30–32]. More generally, trophallaxis acts as a “social glue” mechanism central to group cohesion in socially advanced insect societies [33], reflecting individual- and group-level tendencies toward reciprocal social interaction. In addition, variation in the duration of trophallaxis interaction in bees and face-to-face interactions in humans fall along similar heavy-tailed distributions, suggesting that universal principles may underlie the biology of social interaction networks [34]. Taken together, trophallaxis is a quantifiable analog to other measures of sociability across diverse taxa.

Though automated tracking of individuals engaged in trophallaxis has yielded new insights into the social structure of honey bee society, there are important gaps in the analysis of trophallaxis sociability. These include genomics, transcriptomics, and movement dynamics associated with trophallaxis within the colony. Addressing these gaps promises to reveal deeply conserved neurobiological mechanisms that shape the social brain [28] as well as behavioral mechanisms associated with social bond formation, as in humans [35,36].

In honey bees, transcriptomic analyses have revealed provocative associations between variation in brain gene expression and variation in sociality. Shpigler et al. [20] found significant enrichment for Autism-related genes in the brains of individual honey bees that were consistently unresponsive to task-related social stimuli. In a second study, these “socially unresponsive” bees also were found to have fewer and briefer trophallaxis interactions compared to age-matched individuals that were more responsive to task-related social stimuli [5]. These results established a link between variation in task performance and variation in sociability.

Here we report on behavioral, genomic, transcriptomic, and comparative genomic analyses of trophallaxis sociability. First, we employed automated tracking of individually barcoded bees in colonies to reveal individual variation in this behavior. Second, we quantified movement kinematics to disentangle trophallaxis engagement frequency from overall activity rate and test hypotheses regarding how individual bees influence one another in terms of movement within the hive. Third, we followed these analyses with lab-based behavioral assays that robustly indicate task specialization. We then performed whole-genome resequencing and brain transcriptomic profiling on these extensively profiled individuals to identify genetic variants and brain gene expression profiles associated with variation in behavior (Fig. 1). We also localized the expression in the brain of one particularly interesting gene, *neuroligin-2* (*nlg2*), identified in both genetic and transcriptomic analyses. Finally, using comparative genomics, we explored similarities and differences in variation in trophallaxis sociability, reward processing, and aggression in honey bees and across species as well.

**Figure 1.**
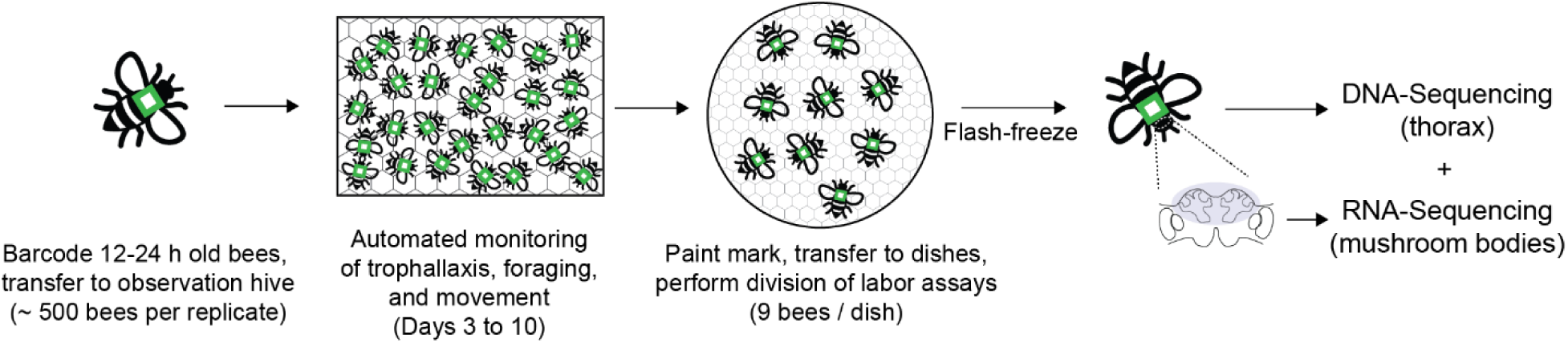
**Experimental design**. For each replicate trial, a mix of ∼500 bees from two source colonies were barcoded and transferred to a single honeycomb frame for automated monitoring of trophallaxis, foraging, and movement behaviors. After 10 days, bees were transferred to a petri dish and subjected to aggression and affiliative care assays. Immediately following behavioral assays, bees were flash-frozen and stored until the thorax and brain were dissected for DNA- and RNA-Sequencing, respectively. Our final sample sizes for whole-genome resequencing and mushroom body RNA-Sequencing were 357 and 176, respectively, and a total of three source colonies were used across two replicate trials.

## Methods

### Animals

All molecular and bioinformatic analyses were performed on bees collected as part of a previous study that addressed different questions [5]. The field- and lab-based behavioral experiments took place at the University of Illinois Bee Research Facility, Urbana, IL, USA, from August to September 2017. Briefly, we used adult worker bees collected from three colonies, each headed by a queen instrumentally inseminated with semen from a single male (drone, “SDI queen”); individuals derived from these matings have an average coefficient of relatedness of 0.75 due to haplodiploidy. To obtain 0- to 24-h-old (one-day-old) bees for age-matching, honeycomb frames containing late-stage pupae were removed from colonies, maintained in a dark incubator at 34°C and 50% relative humidity (RH), and ∼500 individuals were gently swept off the frames the morning of each behavioral experiment. All colonies were filmed when the bees were 3 to 10 days old. On Day 10 of the recording, bees were removed and prepared for laboratory-based behavioral assays, as described below.

### Automated behavioral tracking, image processing, and trophallaxis detection

Single-sided glass-walled observation hives, each containing a colony with barcoded bees, were set up as previously described [5,27,30,31]. Individual bees were cold-anesthetized and a barcode was applied to the dorsal thorax using a small amount of cyanoacrylate glue. Barcoded bees were then gently transferred to an observation hive containing one honeycomb frame provisioned with honey (top 18 rows, ∼200 mg per cell, ∼330 g in total) and pollen paste (next six rows, ∼100 mg per cell, ∼45 g total; pollen paste was made from 45% honey, 45% pollen and 10% water). A naturally mated queen, unrelated to the workers, was lightly anesthetized with carbon dioxide, barcoded, and allowed to recover surrounded by workers in the observation hive.

Observation hives were then transferred to a dark room connected to the outside by a foraging port that was kept closed for the first two days of the experiment to prevent young bees with undeveloped flight muscles from exiting the colony. For recording, we used LED lights (Smart Vision Lights, Muskegon, MI, USA) to illuminate the honeycomb with infrared light, which the bees cannot see. Whole-colony images were captured at a rate of 1 image/s using a Prosilic GT6600 machine vision camera (Allied Vision, Exton, PA). To ensure sharp images, the observation hive glass window was gently replaced daily with a clean one without disturbance to the colony.

Hive images were processed as previously described [5]. Briefly, images were resized and sharpened to improve barcode detection rate, and each bee’s location and orientation were inferred using custom software [27], which also filtered any barcode that was unreadable or resulting from a spurious detection. We then used a convolutional neural network to identify all trophallaxis events by recording when two bees were close (1.7 – 7.4 mm apart), facing each other, and one bee had inserted her extended proboscis (“tongue”) between the open mouthparts of another bee [5,37]. These “raw” trophallaxis events were merged if they were consecutive and <1 min apart, and interactions lasting <3 s or >3 min were removed as spurious detections.

### Bee kinematics quantification

We quantified bee movement as the instantaneous linear (locomotion) and angular (turning) speeds of individuals. Linear speed was defined as the Euclidean distance between a bee’s barcode center at time points t and t+1, divided by the image rate r. Angular speed a was defined as the unsigned angle between a bee’s barcode orientation vector at t and t+1, divided by r. Mean speeds were calculated by averaging these quantities across time. If a bee was not detected at time point t, we recorded no speed measurements for the image captured at time point t+1. We also recorded no speed measurement if a bee moved less than the width of a honeycomb cell (4.9 mm) and turned less than from one cell corner to the next (60°). These thresholds were applied to differentiate actual locomotion from the small movements that a bee may perform while she is stationary (e.g., during allogrooming and other behaviors).

### Social influence analysis

To measure the extent to which the movement of one bee influenced the movement of another bee, we computed the information partial directed coherence (iPDC) [38], employing the AsympPDC package https://www.lcs.poli.usp.br/~baccala/pdc/CRCBrainConnectivity/ [39]. iPDC measures the directed coherence between two or more time-series, enabling estimation of the bidirectional transfer of movement influence-related information. As previously done [5], we focused on the last two days of the recording and limited analysis to periods characterized by proximal interactions (i.e., when two bees were physically close to each other for a minimum amount of time), since it is unlikely that two bees located far apart in the hive would influence each other’s movement directly.

To identify periods of proximal interaction, we computed for each image the Euclidean distance *R* between a focal bee and all other bees, using the *X* and *Y* coordinates of their barcodes. Given a focal bee, we defined *neighbor* as another bee whose distance *R* to the focal bee was equal or lower than a radius *R_max_* for a duration *L* equal or higher than a minimum duration *L_min_* (*R*<=*R_max_, L>*=*L_min_*). We chose *R_max_* = 2cm for being approximately twice the length of a bee’s body, and because previous studies used this value as a measure of proximal interaction in honey bees (Wild et al., 2021). We chose *L_min_* = 30 images as a minimum duration of the proximity interaction in order to have sufficient time-series samples for computing iPDC. In addition, since our aim was to measure social influence during movement, we excluded images where either one of the two bees’ speed was very low as well as images where the two bees were engaging in trophallaxis, during which animals are relatively still. If a focal bee interacted with the same neighboring bee multiple times satisfying the above criteria, we kept data from the first interaction only. In this manner, the identity of the neighbor varied with each proximity interaction. The number of proximity interactions varied across focal bees, depending on the number of unique neighbors they encountered, which could be from the same or different source colony.

We retrieved bees’ *X* and *Y* coordinates during each proximity interaction and built an input dataset to which iPDC was applied. In the dataset, each interaction is represented as a bivariate time-series of at least 30 time points (30 sec), where the first time-series is the focal bee’s *X* or *Y* coordinate and the second time-series is its neighbor’s *X* or *Y* coordinate, respectively. Next, we fit a vector autoregressive model (VAR) to the time-series in each interaction and computed from it the iPDC spectra, from the focal bee to the neighbor and vice versa. iPDC spectra were then averaged across interactions to obtain a single iPDC spectrum for each focal bee, in both directions. Finally, the averaged iPDC spectra were integrated across all frequencies into information flow (*I_flow_*), as adapted from equation 8 in (Takahashi et al., 2010):

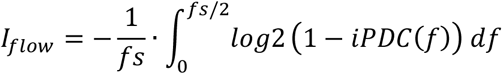

where *fs* is the sampling rate.

We thus obtained a scalar value representing the magnitude of the influence between the focal bee and its neighbors in units of information transfer (bits).

### Behavioral state identification via laboratory-based assays and foraging detection

Behavioral states related to natural colony division of labor (guards, nurses, foragers, generalists, and non-responders) were studied under controlled laboratory conditions as previously described [5]. Briefly, at the conclusion of the recording portion of the experiment, the glass observation window covering the hive was replaced with a Plexiglas window containing resealable portholes, and groups of 9 bees were gently removed, paint-marked with a unique color applied to the dorsal thorax (Testors Paint, Rockford, IL), and transferred to a vertically oriented 100 x 200 mm Petri dish (Thermo Fisher Scientific, Waltham, MA). Dishes contained a tube of honey (∼1.2 ml), 50% sucrose solution (2 ml), and a pollen ball (70% pollen, 30% sucrose solution described above). Bees were given at least 60 min to acclimate to normal fluorescent lighting prior to the start of the behavioral assays.

Established laboratory assays were used to identify aggression, which reflects guarding behavior [40,41] and affiliative caregiving, which reflects nursing behavior [42,43]. Aggression was measured by subjecting groups of bees to a 5-min interaction with a foreign bee (“intruder”) and quantitating aggressive biting and stinging interactions. Affiliative caregiving was measured by exposing groups of bees to a larva for 5 min and quantitating observations of caregiving interactions like food provisioning and inspection. Each group was exposed to both stimuli, and “guards” and “nurses” were identified based on consistent (≥20–30 s) observations of biting and stinging or larval feeding, respectively. We also noted instances of fanning, wax-building, and vibration-signaling behaviors (following descriptions in [44]) to give a more comprehensive depiction of each individual’s behavioral state.

Affiliative care assays were performed in a controlled environment that mimicked the interior of the hive (34°C and 50% relative humidity) under ambient lighting. Aggression assays were performed in a cooler room (28°C) to approximate outdoor temperatures at the hive entrance, where defensive aggression by guards against intruders is most likely to occur. Each dish of 9 bees was tested in both assays in a random order with the second assay performed 60 min after the first. Immediately following the second assay, all bees were flash-frozen in liquid nitrogen, their barcodes removed for identification purposes, and stored at -80°C prior to DNA and RNA analysis.

Because the barcoded colonies were initially composed only of one-day-old bees, precocious foraging occurred during the observation period from days 3-10 [45]. Foragers were identified via automated monitoring of flight in and out of the hive with an entrance monitor attached to the outer portion of the entrance tube connected to the observation hive [30]. Though foraging typically occurs toward the end of a honey bee’s lifetime, in colonies such as these that lack older bees, a subset will exhibit “precocious” foraging to serve the colony’s nutritional needs [46,47]. The entrance monitor contained a small enclosure with a simple maze for slowing bees down to facilitate image capture. A Raspberry Pi Camera Module v1.3 was mounted above the enclosure and separated from the maze by a removable glass window. The camera recorded .mjpg videos at a temporal resolution of 3 images/s from the hours of 07:00 to 19:00 h CST, automatically adjusting for changes in light conditions throughout the day.

Videos of bee activity were first converted to still images using ffmpeg (https://ffmpeg.org/) and then processed to detect barcoded bees, as previously described [31]. Raw incoming and outgoing “passes” were merged by vectoring the bee’s displacement toward or away from the hive entrance. To be labeled a forager, we required bees to be detected taking at least six trips, with more than three trips per day, for any two days of the experiment, and at least 25% of these trips had to be made during peak foraging hours (10:00-15:00 h CST). This is consistent with honey bee foraging behavior under natural conditions [48].

Bees that did not respond to behavioral stimuli during the lab assays and were not observed to forage when in their observation hive were labeled “non-responders” [20]. Bees that showed mild responsiveness (i.e. < 20 s interaction) and displayed some other behaviors (fanning, wax-building, and vibration-signaling behaviors) were referred to as “baseline” bees for which the behavioral state could not be robustly identified but were clearly responsive to stimuli. Bees that exhibited two or more behaviors (i.e., brood care, aggression, and/or foraging) were “generalists.”

### DNA- and RNA-Sequencing and analysis

All DNA- and RNA-Sequencing data were newly generated for this study from the individuals analyzed behaviorally as described above.

#### DNA-Sequencing

DNA was extracted from the thorax of 391 individual bees using the Gentra Puregene Tissue Kit (QIAGEN, Germantown, MD, USA) as previously described (Avalos et al., 2020). Shotgun genomic libraries were prepared using the Illumina DNA (Nextera Flex) sample preparation kit, pooled, quantitated via qPCR, and sequenced on one S4 lane for 151 cycles from both ends of the fragments on an Illumina NovaSeq 6000. Raw fastq files were generated and demultiplexed using the bcl2fastq v2.2 conversion software (Illumina).

#### Mushroom body RNA-Sequencing

We sequenced RNA from 176 individuals, a subset of the larger set used for DNA sequencing. These individuals were selected because they each had a definable behavioral state other than “baseline,” following our above-described assignment strategy. We did not sequence “baseline” bees, as these individuals did not forage or robustly engage social stimuli in the dish assays, but showed higher levels of engagement than non-responders, in addition to other well-known natural behaviors like wax-building or fanning. As we could not reliably characterize these individuals in behavioral terms, molecular profiling would not be as meaningful as for other groups.

Mushroom body (MB) dissection and RNA extraction were performed as previously described [5,49]. We chipped a small window on the anterior surface (frons) of flash-frozen bee heads immersed in a dry/ice ethanol bath before submerging the entire head in RNAlater-ICE Frozen Tissue Transition Solution (Thermo Fisher Scientific) overnight at 20°C. The brain was then fully dissected on wet ice and the MB was removed and stored at −80°C until RNA extraction.

We used the PicoPure RNA Isolation Kit (Thermo Fisher Scientific, Waltham, MA) with DNase treatment (Invitrogen, Carlsbad, CA) as has been previously described [5,20] and quantified via QuBit fluorometer (Thermo Fisher Scientific). RNA-Seq libraries were prepared with the Kapa Hyper Stranded mRNASeq Sample Prep Kit (Illumina, San Diego, CA); libraries were pooled, quantitated via qPCR, and sequenced on an S4 lane for 151 cycles from both ends of the fragment on an Illumina NovaSeq 6000.

### Bioinformatic analyses

#### DNA alignment, variant calling and Genome-Wide Association Study (GWAS)

Whole-genome resequencing yielded at least 40,000,000 paired-end reads per sample (mean [*µ*] ± standard deviation: *µ* = 64,900,620 ± 9,035,972). After adaptor trimming, raw reads were aligned to the honey bee genome, Amel_HAv3.1 (Wallberg et al., 2019), using the Burrows-Wheeler algorithm, and resulting alignment files were sorted and deduplicated. The Samtools “flagstat” report indicated > 99% mapping rates for samples to the honey bee genome, and deduplication of sequenced reads suggested a duplication rate of < 20% (*µ* = 16.8 ± 0.012%). Thus, each sample was sequenced at an average depth of 30x, indicating a high-quality dataset suitable for GWAS.

Variant calling was performed using the Sentieon DNAseq workflow (https://support.sentieon.com/manual/DNAseq_usage/dnaseq/). Following indel realignment and variant calling, which resulted in a VCF file for each of our samples, genotyping was conducted across all samples to generate a single multi-sample VCF file. We then removed indels, multiallelic variants, and variants localized to unplaced scaffolds or mitochondrial genes. The remaining biallelic SNPs were filtered using joint quality measures and overall representation in the dataset, utilizing missing and low-coverage calls in the multi-sample VCF file. No sample showed excessive missing reads. Sequenced genomes were then phased and imputed with Beagle (v5.1) [50,51]. Linkage disequilibrium (LD) pruning was performed using the snpgdsLDpruning command in the R package SNPRelate [52] using the D’ metric with an LD threshold set to 0.3 and mean allele frequency set to 0.2. The resulting ∼2.5 million SNPs were used for downstream analyses.

We derived a genetic relatedness matrix (GRM) using the pcrelate function in GENESIS [53] to account for variation based on kinship, as has been previously done in honey bees [54]. Bees that did not cluster with their colony of origin, and/or bees that were later found to have unreadable barcodes (and therefore unreliable identifying information), were excluded from analysis, leaving us with a final sample size of 357 individuals for DNA analysis.

For individual-level phenotypic association, we utilized the quasi-likelihood approximation in GENESIS, which scores and assigns a *P*-value to each SNP, thus predicting the relevance of the association between a given SNP and phenotype for each genotype. We used GENESIS to identify SNPs significantly associated with trophallaxis sociability or kinematic metrics like locomotion and distance traveled, all averaged over the final two days of the recording experiment.

#### Brain mushroom body (MB) RNA-Sequencing alignments and detection of differentially expressed genes

Each sample yielded at least 13,000 paired-end reads (*µ* = 17,836,571 ± 2,286,238). After demultiplexing, raw reads for each sample were aligned to the honey bee genome, Amel_HAv3.1 [55] using STAR v2.7.3a [56] and reads were counted using the *featureCounts* function in the Subread v2.0 package [57]. We identified the presence of Deformed Wing Virus (DWV), a virus that is naturally widespread in honey bee apiaries and commonly found in the brain [41,58] by also aligning raw reads to the DWV genome. All linear models for identifying differentially expressed genes (DEGs) included percentage of total reads aligned to the DWV genome as a covariate, as well as colony of origin, in order to control for unwanted variation due to natural viral presence or latent genetic variation across colonies. We observed a unique mapping rate of 84.5 ± 0.0845% against the honey bee genome, and an average of 65.7 ± 0.066% of these reads were uniquely mapped to a gene model.

We first identified DEGs associated with sociability by regressing MB gene expression using trophallaxis sociability, averaged over the last two days of the experiment [5], as a continuous predictor. Next, we calculated DEGs associated with each behavioral state (generalist, forager, guard, nurse, or non-responder) by comparing each state to expression levels averaged across the remaining four groups. Each DEG list was generated via the Wald test and corrected for multiple testing using the Benjamini-Hochberg method, FDR < 0.05. Gene Ontology enrichment analysis was performed by converting honey bee gene identifiers to one-to-one orthologs in *Drosophila melanogaster* via a previously published reciprocal best-hit BLAST [58], detecting term enrichment in GOrilla [59], and visualizing results in GO-Figure [60]. Gene list overlap analyses were performed using the GeneOverlap package (https://bioconductor.org/packages/GeneOverlap) with Bonferroni corrections for multiple testing.

#### Spatial analysis of neuroligin-2 gene expression

*In situ* hybridization was performed in January 2023 to provide additional information on one of the genes identified in DNA and RNA analyses that emerged as a particularly strong sociability-associated candidate gene, *neuroligin 2* (*nlg2)*. One-day-old individuals were collected from a honeycomb frame containing brood from a colony kept indoors at the Bee Research Facility. They were kept in a small Plexiglas cage at 35°C and 50% until whole-brain dissections were performed at 7 or 14 days of age.

We followed an established protocol (https://www.protocols.io/view/hcr-rna-fish-protocol-for-the-whole-mount-brains-o-bzh5p386) with minor modifications. Brains were dissected in phosphate-buffered saline (PBS) and fixed in 4% paraformaldehyde (in 1X PBS with Triton-X100; PBS-T) overnight at 4°C. Fixed brains were washed three times in PBS with Tween (Fisher Scientific), dehydrated in serial methanol/PBST washes (25%, 50%, 75% and 100%), and stored at -20°C until use.

Brains were rehydrated in the reverse methanol/PBST serial washes (75%, 50%, 25%) and washed in PBS-T. Brains were pre-hybridized in hybridization buffer (Molecular Instruments, Los Angeles, CA) without probes for 1 h at 37°C. Next, we combined 4 μL of 1 μM HCR detection probes targeting the honey bee *nlg2* gene (Molecular Instruments) with 250 μL hybridization buffer, and incubated each brain in this solution overnight at 37°C. The following day, brains were washed in preheated probe wash buffer (Molecular Instruments) 5 times, 10 min per wash, at 37°C, and then in 5X saline-sodium citrate twice for 5 min per wash. Washed brains were pre-amplified in amplification buffer (Molecular Instruments). Signal amplification hairpin oligos (B1-H1-488 and B1-H2488, from Molecular Instruments) were first denatured at 95°C for 90 s and annealed by cooling on wet ice for 30 min. Amplification hairpins were used at a concentration of 6 μL per 100 μL amplification buffer. Brains were incubated in the amplification probe solutions overnight, followed by two washes in probe wash buffer containing DAPI (Fisher Scientific, R37606). The samples were washed in PBS and cleared with RapiClear® 1.49 (SunJin Lab Co, Hsinchu City, Taiwan) using an established protocol (Bekkouche et al., 2020).

Confocal imaging was performed with a Zeiss LSM 900 inverted confocal microscope (Carl Zeiss, Oberkochen, Germany) with 405 and 488 nm excitation wavelengths. The 20X /0.8, 25X/0.8 objective was used for scanning the whole brain in the tile-scan mode. The Plan-Apochromat 25X/0.8 and 63X/1.4 oil objectives were used for scanning MB sub-regions. Images were viewed and processed in Zeiss ZEN

## Results

### Genome-wide association study (GWAS) on trophallaxis sociability and movement kinematics

We first asked if there are genomic correlates of trophallaxis sociability, utilizing automated tracking of trophallaxis interactions performed across two replicate trials with experimental colonies composed of bees from three source colonies: “R2,” “R8,” and “R41.” Colony 1 was composed of bees from R2 and R41 and Colony 2 was composed of bees from R2 and R8. We generated a ∼2.5 million SNP-set from whole-genome sequencing of 357 bees that we then correlated with trophallaxis sociability. Trophallaxis sociability was quantified as the average number of trophallaxis interactions per bee across the last two days of recording, as in [5].

This analysis identified 18 single nucleotide polymorphisms (SNPs) localized across nine chromosomes (Bonferroni-corrected *P* < 0.05; Fig. 2). Eleven of these SNPs localized to introns of honey bee gene models [61], one localized to a predicted noncoding RNA (LOC113219153), and the remaining six were found outside of genes (Fig. 2; Supp. Table S1). Among the 11 intronic SNPs, two were found in the gene *neuroligin-2*, two in *glutamate receptor ionotropic, NMDA 2B,* which had robust one-to-one orthology with the *Drosophila melanogaster* genes *neuroligin-2* and *NMDA receptor 2*, respectively. A single SNP was found in the following genes, with robust one-to-one orthologs in *D. melanogaster* given in parentheses: *Angiogenic factor with G patch and FHA domains 1* (CG8079), *CCR4-NOT transcription complex subunit 6-like* (no hit in *D. melanogaster*), *nephrin* (*sidestep VIII*), *discoidin domain-containing receptor 2* (no hit in *D. melanogaster*)*, nuclear factor 1 X-type* (*nuclear factor 1*) and uncharacterized genes LOC102655706 (*tartan*) and LOC100577268 (no hit in *D. melanogaster*).

**Figure 2.**
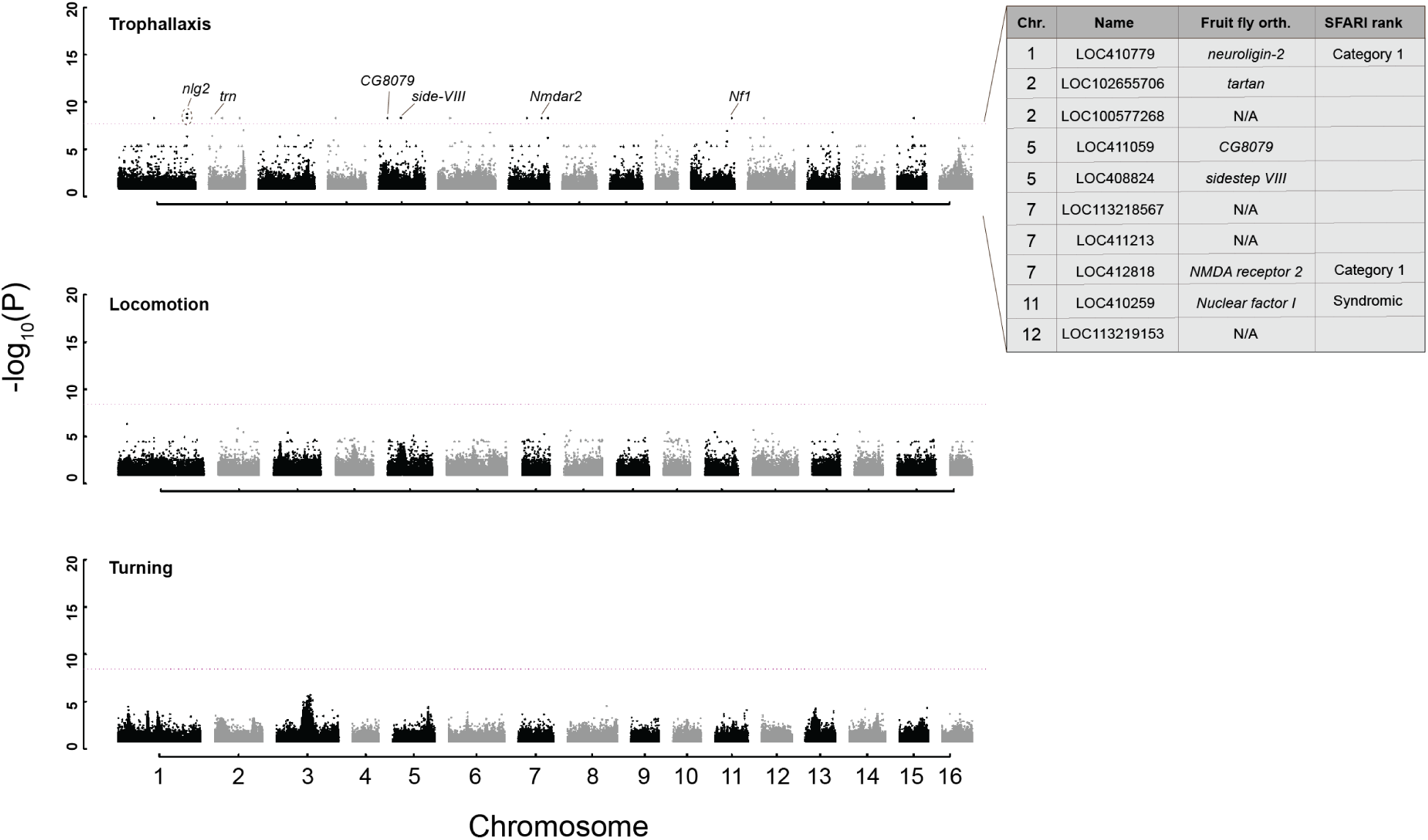
Genomic correlates of honey bee trophallaxis sociability. (Top) Genome-wide association study (GWAS) identified 17 single nucleotide polymorphisms (SNPs) significantly associated with variation in trophallaxis (food-sharing) interactions, 12 of which were localized to the introns of gene models in the honey bee genome (HAv3.1). In vertebrates, variation in the expression or structure of three of these genes, *neuroligin-2*, *NMDA receptor 2*, and *Nuclear factor I* is associated with autism per the Simons Foundation Autism Research Initiative (SFARI) database. (Middle and bottom) In contrast, we did not identify any SNPs associated with variation in locomotion or turning kinematics, movements that contribute to, but do not fully encompass, trophallaxis interactions. Dashed red line indicates significance threshold (Bonferroni-corrected *P*-value < 0.05).

In a comparative bioinformatic analysis, n*euroligin-2* (*nlg2*) and *NMDA receptor 2* (*nmdar2*) were found to be Category 1 (“high confidence”) genes in the Simons Foundation Autism Research Initiative (SFARI) gene database (https://gene.sfari.org/), meaning they have been clearly implicated in human Autism Spectrum Disorder. Similarly, *nuclear factor 1 X-type* (*Nf1*) is characterized as a “Syndromic” autism gene, meaning its dysfunction is associated with the development of autistic traits that are not typically associated with a conventional autism diagnosis.

We performed the same GWAS analysis for two other behaviors: locomotion and turning. Like trophallaxis, these behaviors require fine motor control and therefore contribute to trophallaxis interactions, but without obvious social components. (Fig. 2). In contrast to our findings for trophallaxis sociability, GWAS for these two behaviors did not result in the detection of significantly associated SNPs.

Neuroligin-2 (nlg2*) and colony sociability*

Of the three source colonies assayed in this study, individuals from source colony R41 were found to all be homozygous for the alternative allele (i.e., containing alleles that were different from the reference locus in the honey bee genome) for the majority of identified SNPs (Fig. 3a; Supp. Fig. S1). In contrast, individuals from source colonies R2 and R8 were predominantly heterozygous for the more prevalent reference allele. This result prompted us to investigate the genetics of colony-level variation in behavior.

**Figure 3.**
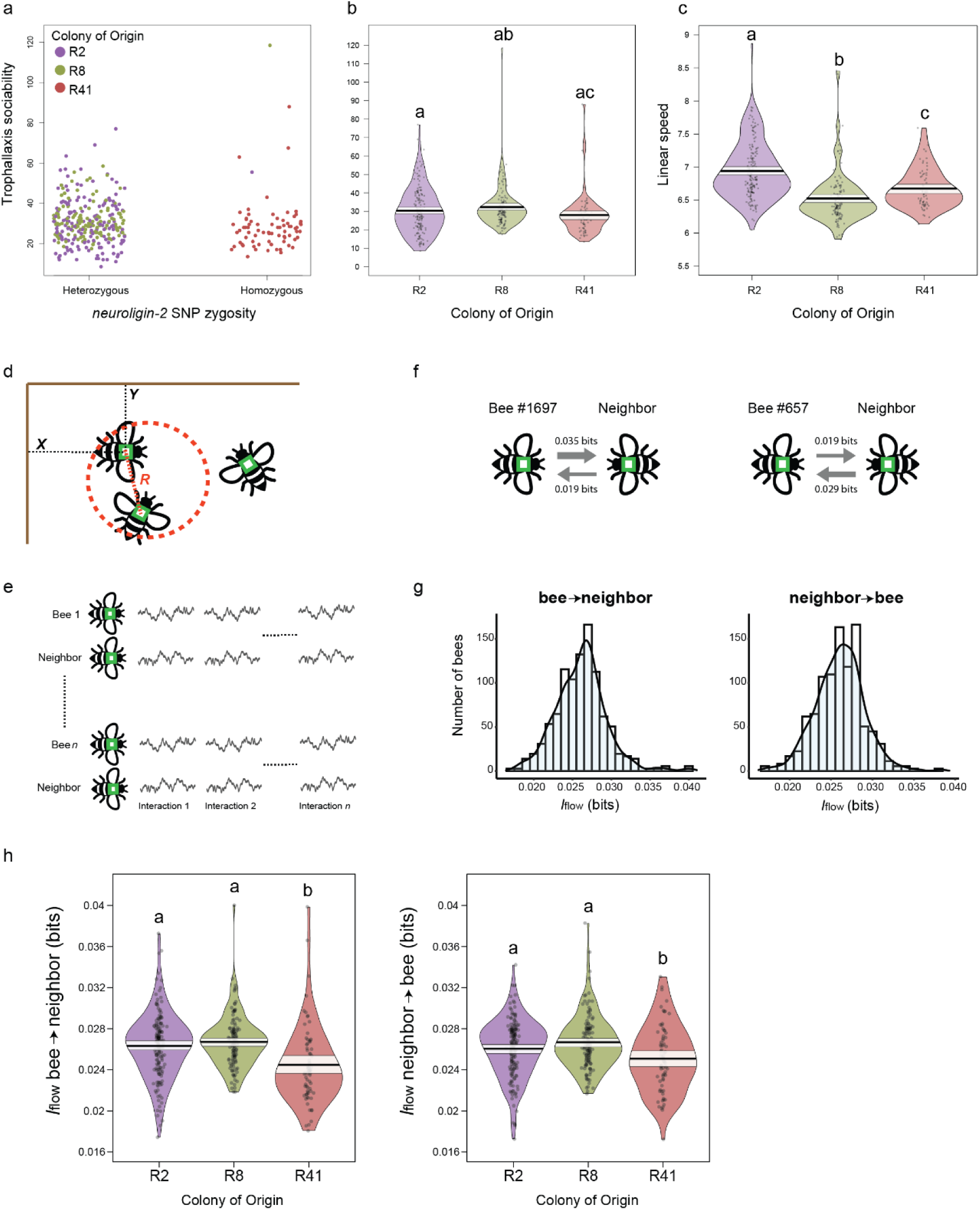
Genotype-specific variation in motor synchrony. (a) A single source colony, “R41,” was found to be homozygous for *nlg2* as well as the majority of trophallaxis-associated SNPs, while other colonies in this experiment contained the more prevalent reference allele. (b) Colony “R41” also demonstrated the lowest levels of trophallaxis sociability, yet (c) was not the slowest-moving colony. (d) Images of proximal interaction were selected where two bees were physically close to each other (*R* ≤ 2 cm, where *R* is the Euclidean distance between the two bees’ barcodes). The *X* and *Y* coordinates of the barcode represent a given bee’s position in Euclidian space. (e) Drawing representing the dataset to which iPDC was applied. Each proximal interaction (interaction 1, … interaction *n*) is represented as a bivariate time-series of at least 30 time points (30 sec), where the first time-series is the focal bee’s *X* or *Y* coordinate and the second time-series is its neighbor’s *X or Y* coordinate, respectively. For each focal bee, the identity of the neighbor varied across interactions. iPDC was calculated within each paired time-series and averaged across interactions to subsequently compute information flow in both directions. (f) Example of two bees with different levels of social influence. Arrow thickness is proportional to information transfer. On average, Bee #1697 influenced the movement of its neighbor with *I_flow bee->neighbor_* = 0.035 bits, and was influenced by the movement of its neighbor with *I_flow neighbor->bee_* = 0.019 bits. (g) Distributions of information flow across bees, representing the magnitude of movement influence-related information. (h) As for trophallaxis sociability, “R41” bees were also least likely to influence, or be influenced by, other bees in their experimental cohort in terms of motor synchrony. Violin plots are constructed as follows: points represent raw data, solid black lines represent the mean, pale white shading above and below the mean represent a 95% confidence interval, and plot shape represents a smoothed density curve outlining the distribution of raw data. Letters above violin plots represent significance (*P*-value < 0.05) from a between- group Tukey Post-Hoc analysis following a one-way ANOVA.

Trophallaxis sociability significantly varied with source colony (ANOVA: F_(2,354)_ = 3.165, *P* = 4.24e-16), and bees from source colony R41 displayed significantly lower trophallaxis rates than those from source colony R8 (Tukey HSD post-hoc test, *P* = 0.034) but not R2 (*P* = 0.23; Fig. 3b). In addition, R2 individuals were significantly less social when co-housed with R41 individuals in Colony 1 than with R8 individuals in Colony 2 (Wilcoxon rank-sum test: W = 6833.5, *P* = 4.17e-11), further implicating R41 as significantly less sociable than the other genotypes.

While individuals from source colony R41 demonstrated the lowest levels of trophallaxis sociability, they were not the slowest moving relative to bees from the other two source colonies. Locomotion was significantly associated with source colony (ANOVA: F_(2,349)_ = 12.52, *P* < 4.24e-16), and each colony significantly differed from one another in this measurement (Tukey HSD post-hoc test, *P* < 0.005 for all comparisons). However, movement dynamics could not explain the observed variation in trophallaxis rate (Fig. 3c).

### Social influences on movement in the hive

To explore bee social interactions further, we measured the extent to which a given bee’s movement influenced, and was influenced by, the movement of a nestmate. We implemented information Partial Directed Coherence (iPDC), which allows for the measuring of bidirectional influences between signals [38]. To this end, we identified periods of proximal interaction (i.e., when two bees were physically close to each other), retrieved the *X* and *Y* coordinates of each bee relative to the upper-left corner of the hive, and quantified iPDC from the coordinate time-series of one bee to the coordinate time-series of its neighbor and vice versa. From iPDC, we then computed information flow (*I_flow_*) (Fig. 3d-f; see also Methods). Higher *I_flow_* values indicate higher information transfer from the past location of a bee to the current location of its neighbor, and we used this metric as measure of social influence. *I_flow_* was approximately normally distributed across bees (Fig. 3g; *I_flow bee->neighbour_* µ =0.0261 bits, σ=0.0033; *I_flow neighbour->bee_* µ =0.0261 bits, σ=0.0029) and the values observed are consistent with those reported for other animal species when assessing social influence in movement dynamics [62]. We then asked whether bees of different colonies exhibited different levels of social influence. Information flow significantly varied with source colony (Fig. 3h; Supp. Fig. S2; one-way ANOVA *I_flow bee->neighbour_*: F_(2,352)_ = 12.26, *P* = 7.11e-06; *I_flow neighbour->bee_*: F_(2,352)_ = 7.1, *P* = 9.51e-4). Post-hoc comparisons revealed no difference between colonies R2 and R8 (Tukey HSD *I_flow bee->neighbour_*: *P* = 0.58; *I_flow neighbour->bee_*: *P* = 0.17). In contrast, consistent with the trophallaxis results, bidirectional *I_flow_* was lowest for individuals from colony R41 compared to R2 (Tukey HSD *I_flow bee->neighbour_*: *P* = 1.37e-4; *I_flow neighbour->bee_*: *P* = 0.045) and R8 (Tukey HSD *I_flow bee->neighbour_*: *P* = 7.6e-6; *I_flow neighbour->bee_*: *P* = 5.69e-4), suggesting that R41 bees show reduced influence on, and from, their neighbor’s movement during time of social proximity.

### Mushroom body (MB) transcriptomic profiles and trophallaxis sociability

We next asked how trophallaxis sociability is related to gene expression, focusing on the mushroom bodies (MB), a higher-order center of sensory integration, decision-making [14], and social regulation [42]. Unlike GWAS, transcriptome profiling generates arrays of genes that associate with a particular continuous or discrete phenotypic variable. Deriving lists of genes with expression values correlated with trophallaxis rate, for example, allows us to test for statistically significant overlap between our findings and previous studies of other behaviors, such as reward processing. We used RNA-Sequencing (RNA-Seq) to profile the MB transcriptome of a subset of bees included in the GWAS, selecting individuals that were reliably identified as generalists, foragers, guards, nurses, or non-responders, following criteria from previous studies [5,20,40].

Non-responders were transcriptionally distinct from generalists, foragers, guards, and nurses, as in [20] ; Supp. Tables S2-S6). A GO enrichment performed on genes that were upregulated in all responders compared to non-responders identified only “chemosensory behavior” and “learning or memory” as enriched in responders. For genes upregulated in non-responders, only a number of less specific metabolic terms were identified as enriched (Fig. 4a).

**Figure 4.**
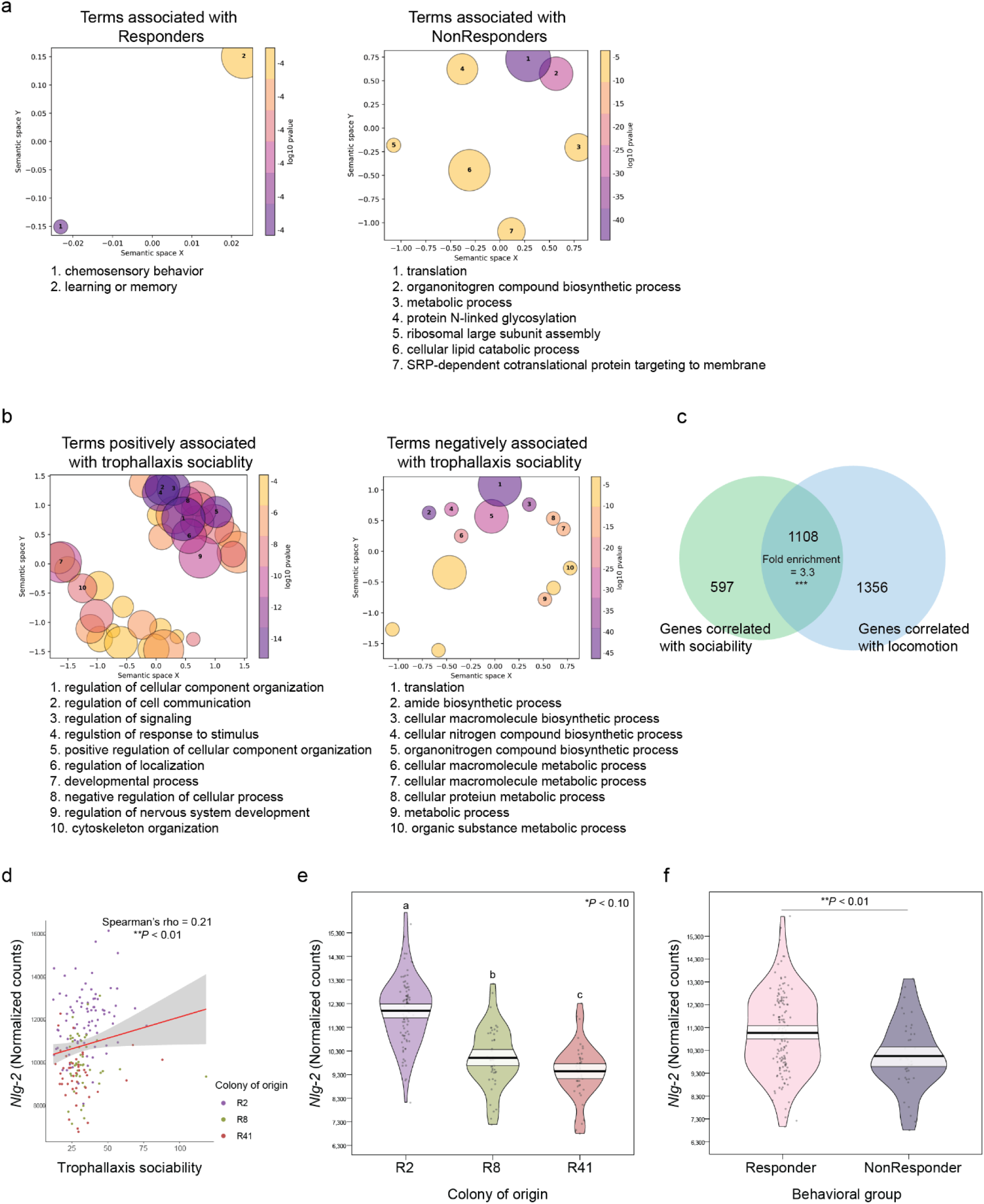
Neuromolecular correlates of trophallaxis sociability and social responsiveness in the honey bee mushroom bodies (MB). (a) Gene Ontology (GO) analysis identified highly specific terms associated with chemosensation and learning enriched in genes upregulated in Responders, while generic metabolic process terms were enriched in genes upregulated in NonResponders. (b) GO enrichment analysis performed on genes positively correlated with trophallaxis frequency revealed the enrichment of several terms related to cell signaling and stimulus response, while genes negatively associated with trophallaxis were more strongly related to biosynthesis and metabolism, and (c) this gene set was significantly similar to genes with expression levels correlated with locomotion (hypergeometric overlap test). (d) *Nlg2* levels were weakly but significantly correlated with trophallaxis frequency, (e) significantly lowest in colony “R41” (one-way ANOVA with Tukey post-hoc test), and (f) differentially expressed in Responders compared to NonResponders in a lab-based assay examining behavioral responsiveness to social stimuli (Wilcoxon rank-sum test). For GO plots, circle diameter is inversely correlated to term specificity in the GO hierarchy, and more similar terms are more closely clustered in semantic space. Violin plots are constructed as previously described.

We identified 920 and 785 genes with expression values that positively or negatively correlated with individual differences in sociability, respectively (Supp. Table S7). Gene Ontology (GO) enrichment analysis identified distinct terms for each set of genes, with positively associated genes enriched for several regulatory processes in the context of cellular communication and signaling and negatively associated genes enriched for processes associated with translation and macromolecule biosynthesis (Fig. 4b). In contrast to our GWAS results, we also identified a significant overlap of 1108 genes with expression values that correlated both with trophallaxis sociability and locomotion (hypergeometric test, fold enrichment = 3.3, *P* = 2.75e-239; Fig. 4c; Supp. Table S8).

Comparing our data with previously published results, we did not identify a significant overlap of genes positively or negatively associated with trophallaxis sociability and those associated with food reward or anticipation of food reward (hypergeometric test, *P* > 0.10) [63,64].

Neuroligin-2 *RNA-Seq expression and fluorescent* in situ *hybridization*

We compared *nlg2* expression in the MB in non-responders to the average expression in generalists, foragers, guards, and nurses. Expression levels were significantly associated with trophallaxis sociability (Spearman’s rho = 0.21, *P* = 0.0063, Fig. 4d), source colony (Tukey HSD post-hoc test on significant ANOVA [F_(2, 169)_ = 69.762, *P* < 2e-16], *P* < 0.10 for each comparison, Fig. 4e), and overall social responsiveness, when (“responders,” Wilcoxon rank sum test, W = 3682, *P* = 0.0014, Fig. 4f).

Motivated by the association of *nlg2* and trophallaxis sociability in both GWAS (Fig. 2) and RNA-Seq results (Fig 3d-f), we asked where *nlg2* is expressed in the brain, and if its regional localization overlaps with regions known to be associated with social decision-making. We collected bees of different ages and used HCR-FISH to visualize *nlg2* expression. We found *nlg2* localization primarily in neuronal somata and largely absent from synaptically dense neuropil when compared to control samples where no gene-specific probes were used (Fig. 5a-b), as expected. The Class I Kenyon cells of the mushroom bodies (MB) appeared to have higher expression, particularly in the densely packed neurons at the center of the MB calyces (Fig. 5c), and expression was weaker in the surrounding, more loosely packed cells (Fig. 5d). We also observed expression in MB extrinsic neurons and dorsal midline populations of cells that compose the pars intercerebralis (Fig. 5e), a small population of large neurosecretory cells vital for nutritional signaling [65]. Some optic lobe cell populations, especially those most proximal to the central brain, also showed discernable fluorescence indicating transcription of *nlg2*.

**Figure 5.**
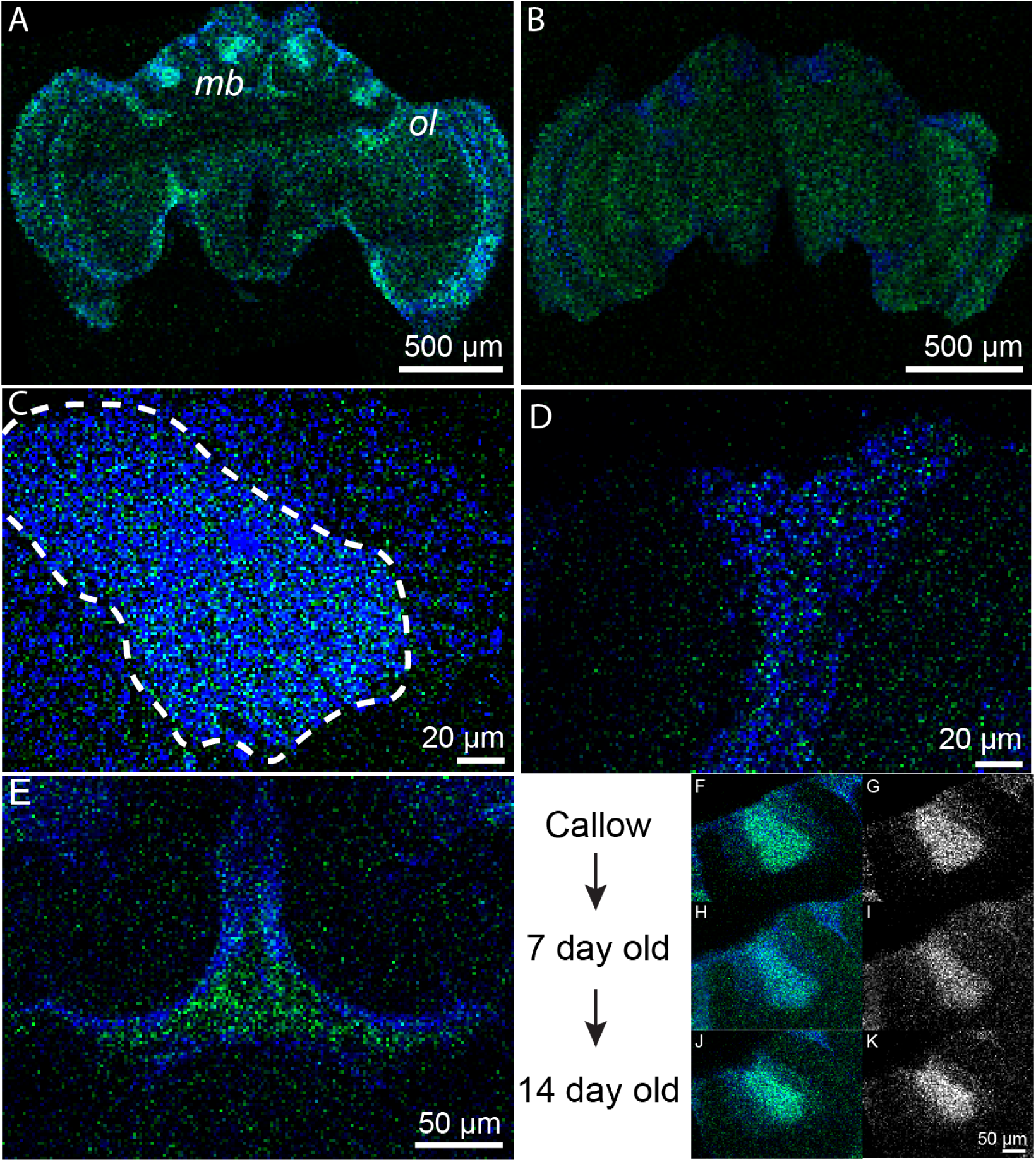
Whole-mount fluorescent *in situ* hybridization (smFISH) reveals *neuroligin-2* (*nlg2*) expression patterns in the developing adult honey bee brain. (a) *Nlg2* is widely expressed in the adult worker brain as detected via HCR probes, while (b) no staining was observed with antisense probes. (c) Expression levels were highest in the mushroom body (MB) Kenyon cells (KC), specifically the inner-compact KC (dashed white line), yet (d) were observed in the outer-compact KC as well. (e) We also identified strong signaling along the dorsal midline of the brain, in or around the neurosecretory cells that compose the pars intercerebralis. (f-k) *Nlg2* expression was apparent throughout adulthood in the worker honey bee brain, though levels appeared highest in callows and two-week-old bees (left and right columns are color and greyscale depictions of the same image, respectively). For each panel, *nlg2* localization is shown in green and nuclei counterstained with DAPI are shown in blue.

As our study considered 10-day-old bees, we sought to qualitatively assess how spatiotemporal expression of *nlg2* varied before and after this age. We performed FISH on one-, seven-, and 14-day-old bees, and found that *nlg2* expression was present in one-day-olds but weak in seven-day-old bees before returning to discernable levels in 14-day-old bees (Fig. 5f-k).

## Discussion

Social proclivities vary within animal societies, driving a continuous distribution of individual tendencies to associate with others. This is true for many species, from humans to honey bees: some individuals pursue regular social contact and maintain large networks while others engage less. Social interactions are multifaceted, as are their drivers, which integrate across physiology, developmental and motivational state, genetics, and past experience. It thus remains challenging to quantify social interactions and their molecular underpinnings. We used a combination of high-throughput behavioral tracking [27,37], laboratory-based behavioral assays [5,42], genome resequencing, brain transcriptomics, and comparative genomics to explore honey bee sociability and its molecular underpinnings. We focused on trophallaxis interactions and inter-individual motor coordination. Our results indicate deep homology in the molecular roots of social behavior shared between honey bees and vertebrates while also revealing divergent mechanisms.

Our genome-wide association study (GWAS) identified 18 single nucleotide polymorphisms (SNPs) associated with variation in trophallaxis sociability. In contrast, no SNPs were identified when a GWAS was performed on locomotion or turning rate, kinematic metrics that contribute to trophallaxis. Like any social interaction, trophallaxis is more than the sum of its parts: rapid movement may support more social interactions but our results indicate it does not alone drive variation in tendency to interact in this way. By standards of human GWAS, our sample size (N = 357 individuals) was small, but similar numbers of sociability-associated SNPs have been identified from GWAS with much larger human cohorts [25], reflecting the complexity of factors underpinning sociability.

Over half of the identified SNPs were localized to the intronic regions of genes. Using the Simons Foundation Autism Research Initiative (SFARI) gene database (https://gene.sfari.org/), we identified *neuroligin-2* (*nlg2*) and *NMDA receptor 2* (*nmdar2*) as Category 1 (“high confidence”) genes, meaning they have been strongly implicated in human autism. Another gene with an intronic SNP, *nuclear factor 1 X-type* (*Nf1*) is characterized as a “Syndromic” autism gene, meaning its dysfunction is associated with the development of autistic traits that are not typically associated with a conventional autism diagnosis. In humans, impaired glutamatergic signaling by way of NMDAR disruption has clear associations with the development of autism [66], while glutamate is best known in insects for its role at the neuromuscular junction [67]. However, NMDAR activity is causally associated with memory formation in honey bees [68], and its function may therefore contribute to the reinforcement of trophallaxis relationships, which ultimately rely on reciprocity between nestmates.

The identification of two *nlg2* SNPs associated with variation in trophallaxis sociability contributes to a comparative framework for understanding the molecular building blocks of social behavior across highly diverged species. Neuroligins are cell surface proteins involved in the formation and remodeling of synapses in the central nervous system in an isoform-specific manner, and *nlg2* acts exclusively at inhibitory synapses [69]. In vertebrates, genomic variation in, or disrupted expression of, *nlg2* or its interaction partners is associated with forms of neurodivergence like autism, schizophrenia, and anxiety, each of which is linked to differences in how individuals associate with their social environment [2,70,71]. In *Drosophila melanogaster*, social behavior also is affected by mutations in *nlg2* [72].

Several correlative findings suggest a role for *nlg2* in honey bee trophallaxis sociability. First, the two intronic variants (SNPs) in the honey bee *nlg2* gene that we identified were predominantly restricted to individuals from one source colony, and those individuals displayed low levels of trophallaxis sociability compared to those from other source colonies. We probed other kinematic aspects of these bees and found that this lower level of sociability was not due to slower movement. Second, *nlg2* expression levels in the mushroom bodies (MB), a higher-order sensory integration center in the bee brain [14,73], were also significantly correlated with trophallaxis rate and social responsiveness across all three tested colonies. Third, a strong signal of *nlg2* expression was localized to the MB and appeared to overlap the pars intercerebralis, a set of neurosecretory cells that mediate the relationship between diet, nutrition, and division of labor [65]. Trophallaxis sociability in honey bees illustrates the importance of food-sharing as a foundation for animal society. We speculate that *nlg2* may have played a role in the shaping of ancestral feeding circuitry to drive stable social network formation in this social bee lineage, as may be the case for other social animals. Consuming but not fully digesting food requires a neurobiological change to bypass the rewarding nature of feeding events, yet such a mechanism provides a foundation for the formation of long-term social bonds [74].

We also explored the synchronization of body movement between pairs of individual bees, some not necessarily engaged in trophallaxis. This model of social influence is similar to trophallaxis in that it requires coordination of one individual’s body with another among honey bees located close to each other, but also directly quantifies a measure of social influence identified in other species [62]. We found that the *ngl2* allele associated with reduced trophallaxis also was associated with reduced motor synchrony between bees. This means that the movement of bees in the colony with high frequencies of this allele had less influence on, and were less influenced by, the movement of other bees. Interpersonal synchrony characterizes temporally or physically aligned behaviors in humans that naturally emerge between individuals in a social group, and such bidirectional motor influence tends to be diminished among autistic individuals and their peers [75,76]. It is interesting to note that casual observations revealed no motor deficits in the colony with high frequencies of this allele and it is therefore unclear if negative fitness consequences exist. It may be that such variation is well-tolerated within the honey bee population in the context of colony performance.

Social interactions help establish and reinforce group membership and, in the context of food-sharing, facilitate mutually rewarding altruistic bonds between individuals [77,78]. In this regard, we predicted trophallaxis to be an intrinsically rewarding behavior, as is honey bee dance communication [63]. However, this prediction was not supported by comparative transcriptomic analysis. We did not identify gene expression similarities between the present study and previous MB transcriptomic characterization of reward processing [63,64]. Perhaps this finding is related to the fact that dopaminergic neurons, which contribute to reward seeking, expectation and acquisition in honey bees [79,80] are sparsely distributed in small clusters throughout the bee brain [81,82], earlier findings corroborated by more recent single-cell brain analyses [83,84]. Perhaps the bulk sequencing of the MB performed in our study both diluted and excluded dopaminergic neurons. It may also be that food donation versus reception, which we did not distinguish in this study, supports different brain gene expression signatures.

While we identified significant variation in social proclivities across and within honey bee colonies, and have started to elucidate its molecular roots, the effects of this variation on colony health and well-being are unknown. Basic evolutionary theory supports the speculation that socially divergent phenotypes are either tolerated or preserved across diverse animal lineages due to supportive, or perhaps even vital, roles they contribute to their society. It could be useful to explore this issue in honey bees, given serious concerns about honey bee health and worldwide colony population declines over the past two decades.

The expression of individual differences in social behavior, from humans to honeybees, is derived from a constellation of interacting neurobiological and physiological factors. While no single gene or genetic variant alone drives sociability, the identification of shared genomic underpinnings of social interactivity across diverse taxa hints at conserved molecular building blocks of social life. We anticipate new tools for automated tracking will continue to improve the granularity of such behavioral comparisons across social animals [28,37], further promoting a role for social insects in elucidating how the brain and genome have evolved to support the intricate social lives observed across diverse animal societies.

## Acknowledgments

The authors thank A. Hernandez and C. Wright at the Roy J. Carver Biotechnology Center (UIUC) for sequencing services and members of the Robinson and Cook labs for valuable comments and feedback that improved the manuscript. This work was supported by the European Union’s Horizon 2020 Research and Innovation Program under ERC-2017-StG Grant Agreement 757583 (Brain2Bee; to JLC and GER). IMT is presently supported by the Lewis-Sigler Institute for Integrative Genomics as a Lewis-Sigler Scholar.

## Supplementary Figures

Supplementary Figure S1. Zygosity for trophallaxis-associated single-nucleotide polymorphisms (SNPs). All SNPs are described in Supplementary Table S1; 17 out of 18 SNPs are shown, with the remaining SNP shown in Fig. 3a. Supplementary Figure S2. Genotype-specific variation in motor synchrony. iPDC was applied to pairs of bees’ *X* and *Y* coordinate time-series, obtaining information flow (*I*flow) between the movement of a bee and its neighbor along the X and Y axis, respectively. *I*flow was then averaged across the two coordinates obtaining a single *I*flow for each bee (Fig. 3h). Violin plots are constructed as follows: points represent raw data, solid black lines represent the mean, pale white shading above and below the mean represent a 95% confidence interval, and plot shape represents a smoothed density curve outlining the distribution of raw data. Letters above violin plots represent significance (*P*-value < 0.05) from a between-group Tukey Post-Hoc analysis following a one-way ANOVA (for *X* coordinate *I_flow bee->neighbour_* : F_(2,352)_ = 8.03, *P* = 3.87e-04, *I_flow neighbour->bee_*: F_(2,352)_ = 7.02, *P* = 0.001); for *Y* coordinate *I_flow bee->neighbour_* : F_(2,352)_ = 6.92, *P* = 0.001, *I_flow neighbour- >bee_*: F_(2,352)_ = 1.4, *P* = 0.248). Tukey HSD for X coordinate *I_flow bee->neighbour_*: *P* = 0.58; *I_flow neighbour->bee_*: *P* = 0.17).

## Supplementary Tables

Supplementary Table S1: Annotations and test statistics for 18 single-nucleotide polymorphisms (SNPs) associated with variation in trophallaxis sociability via genome-wide association study (GWAS).

Supplementary Table S2: Differentially expressed genes detected in non-responders relative to remaining behavioral groups.

Supplementary Table S3: Differentially expressed genes detected in generalists relative to remaining behavioral groups.

Supplementary Table S4: Differentially expressed genes detected in foragers relative to remaining behavioral groups.

Supplementary Table S5: Differentially expressed genes detected in guards relative to remaining behavioral groups.

Supplementary Table S6: Differentially expressed genes detected in nurses relative to remaining behavioral groups.

Supplementary Table S7: Genes with expression levels that correlate with trophallaxis sociability. Supplementary Table S8: Genes with expression levels that correlate with locomotion.

## Notes

### Competing Interest Statement

The authors have declared no competing interest.

### Summary of Updates

Added a new Figure 1 that provides a graphical depiction of the behavioral and molecular experiments performed in this study.

## References

1. Gartland LA, Firth JA, Laskowski KL, Jeanson R, Ioannou CC. Sociability as a personality trait in animals: methods, causes and consequences. Biol Rev. 2022;97: 802–816. doi:10.1111/brv.12823

2. Barak B, Feng G. Neurobiology of social behavior abnormalities in autism and Williams syndrome. Nat Neurosci. 2016;19: 647–655. doi:10.1038/nn.4276

3. Bonnell TR, Vilette C, Henzi SP, Barrett L. Network reaction norms: taking account of network position and plasticity in response to environmental change. Behav Ecol Sociobiol. 2023;77: 35. doi:10.1007/s00265-023-03300-2

4. Friedman DA, Johnson BR, Linksvayer TA. Distributed physiology and the molecular basis of social life in eusocial insects. Horm Behav. 2020;122: 104757. doi:10.1016/j.yhbeh.2020.104757

5. Traniello IM, Hamilton AR, Gernat T, Cash-Ahmed AC, Harwood G, Ray AM, et al. Context-dependent influence of threat on honey bee social network dynamics and brain gene expression. J Exp Biol. 2022;225. doi:10.1242/jeb.243738

6. Taylor SE, Klein LC, Lewis BP, Gruenewald TL, Gurung RAR, Updegraff JA. Biobehavioral responses to stress in females: Tend-and-Befriend, not Fight-or-Flight. Psychol Rev. 2000;107: 411–429. doi:10.1037/0033-295x.107.3.411

7. López-Tobón A, Trattaro S, Testa G. The sociability spectrum: evidence from reciprocal genetic copy number variations. Mol Autism. 2020;11: 50. doi:10.1186/s13229-020-00347-0

8. Fischer EK, O’Connell LA. Hormonal and neural correlates of care in active versus observing poison frog parents. Horm Behav. 2020;120: 104696. doi:10.1016/j.yhbeh.2020.104696

9. Kabelik D, Weitekamp CA, Choudhury SC, Hartline JT, Smith AN, Hofmann HA. Neural activity in the social decision-making network of the brown anole during reproductive and agonistic encounters. Horm Behav. 2018;106: 178–188. doi:10.1016/j.yhbeh.2018.06.013

10. Johnson ZV, Hegarty BE, Gruenhagen GW, Lancaster TJ, McGrath PT, Streelman JT. Cellular profiling of a recently-evolved social behavior in cichlid fishes. Nat Commun. 2023;14: 4891. doi:10.1038/s41467-023-40331-9

11. Newman SW. The medial extended amygdala in male reproductive behavior: A node in the mammalian social behavior network. Ann N York Acad Sci. 1999;877: 242–257. doi:10.1111/j.1749-6632.1999.tb09271.x

12. O’Connell LA, Hofmann HA. The Vertebrate mesolimbic reward system and social behavior network: A comparative synthesis. J Comp Neurol. 2011;519: 3599–3639. doi:10.1002/cne.22735

13. O’Connell LA, Hofmann HA. Evolution of a vertebrate social decision-making network. Science. 2012;336: 1154–1157. doi:10.1126/science.1218889

14. Strausfeld NJ. Arthropod Brains: Evolution, Functional Elegance, and Historical Significance. Harvard University Press; 2012.

15. Wilson EO. The insect societies. Cambridge, MA: Harvard University Press; 1971.

16. Page RE, Robinson GE. The genetics of division of labour in honey bee colonies. Adv Insect Physiol. 1991;23: 117–169. doi:10.1016/s0065-2806(08)60093-4

17. Whitfield CW, Cziko A-M, Robinson GE. Gene expression profiles in the brain predict behavior in individual honey bees. Science. 2003;302: 296–299. doi:10.1126/science.1086807

18. Zayed A, Robinson GE. Understanding the relationship between brain gene expression and social behavior: Lessons from the honey bee. Genetics. 2012;46: 591–615. doi:10.1146/annurev-genet-110711-155517

19. Cook JL, Robinson GE. Comparative genomics and the roots of human behavior. Trends Cogn Sci. 2023;27: 230–232. doi:10.1016/j.tics.2022.12.012

20. Shpigler HY, Saul MC, Corona F, Block L, Ahmed AC, Zhao SD, et al. Deep evolutionary conservation of autism-related genes. Proc Natl Acad Sci. 2017;114: 9653–9658. doi:10.1073/pnas.1708127114

21. Rittschof CC, Bukhari SA, Sloofman LG, Troy JM, Caetano-Anollés D, Cash-Ahmed A, et al. Neuromolecular responses to social challenge: Common mechanisms across mouse, stickleback fish, and honey bee. Proc Natl Acad Sci. 2014;111: 17929–17934. doi:10.1073/pnas.1420369111

22. Traniello IM, Chen Z, Bagchi VA, Robinson GE. Valence of social information is encoded in different subpopulations of mushroom body Kenyon cells in the honeybee brain. Proc R Soc B. 2019;286: 20190901. doi:10.1098/rspb.2019.0901

23. Arechavaleta-Velasco ME, Alcala-Escamilla K, Robles-Rios C, Tsuruda JM, Hunt GJ. Fine-Scale linkage mapping reveals a small set of candidate genes influencing honey bee grooming behavior in response to varroa mites. PLoS ONE. 2012;7: e47269. doi:10.1371/journal.pone.0047269

24. Kocher SD, Mallarino R, Rubin BER, Yu DW, Hoekstra HE, Pierce NE. The genetic basis of a social polymorphism in halictid bees. Nat Commun. 2018;9: 4338. doi:10.1038/s41467-018-06824-8

25. Bralten J, Mota NR, Klemann CJHM, Witte WD, Laing E, Collier DA, et al. Genetic underpinnings of sociability in the general population. Neuropsychopharmacology. 2021;46: 1627–1634. doi:10.1038/s41386-021-01044-z

26. Gal A, Saragosti J, Kronauer DJ. anTraX, a software package for high-throughput video tracking of color-tagged insects. eLife. 2020;9: e58145. doi:10.7554/elife.58145

27. Gernat T, Rao VD, Middendorf M, Dankowicz H, Goldenfeld N, Robinson GE. Automated monitoring of behavior reveals bursty interaction patterns and rapid spreading dynamics in honeybee social networks. Proc Natl Acad Sci. 2018;115: 1433–1438. doi:10.1073/pnas.1713568115

28. Traniello IM, Kocher SD. Integrating computer vision and molecular neurobiology to bridge the gap between behavior and the brain. Curr Opin Insect Sci. 2024;66: 101259. doi:10.1016/j.cois.2024.101259

29. Wolf SW, Ruttenberg DM, Knapp DY, Webb AE, Traniello IM, McKenzie-Smith GC, et al. NAPS: Integrating pose estimation and tag-based tracking. Methods Ecol Evol. 2023. doi:10.1111/2041-210x.14201

30. Geffre AC, Gernat T, Harwood GP, Jones BM, Gysi DM, Hamilton AR, et al. Honey bee virus causes context-dependent changes in host social behavior. Proc Natl Acad Sci. 2020;117: 10406–10413. doi:10.1073/pnas.2002268117

31. Jones BM, Rao VD, Gernat T, Jagla T, Cash-Ahmed AC, Rubin BE, et al. Individual differences in honey bee behavior enabled by plasticity in brain gene regulatory networks. eLife. 2020;9: e62850. doi:10.7554/elife.62850

32. Wild B, Dormagen DM, Zachariae A, Smith ML, Traynor KS, Brockmann D, et al. Social networks predict the life and death of honey bees. Nat Commun. 2021;12: 1110. doi:10.1038/s41467-021-21212-5

33. Wcislo WT. Trophallaxis in weakly social bees (Apoidea). Ecol Èntomol. 2016;41: 37–39. doi:10.1111/een.12289

34. Choi SH, Rao VD, Gernat T, Hamilton AR, Robinson GE, Goldenfeld N. Individual variations lead to universal and cross-species patterns of social behavior. Proc Natl Acad Sci. 2020;117: 31754–31759. doi:10.1073/pnas.2002013117

35. Glass D, Yuill N. Social motor synchrony in autism spectrum conditions: A systematic review. Autism. 2024;28: 1638–1653. doi:10.1177/13623613231213295

36. Glass D, Yuill N. Evidence of mutual non-verbal synchrony in learners with severe learning disability and autism, and their support workers: a motion energy analysis study. Front Integr Neurosci. 2024;18: 1353966. doi:10.3389/fnint.2024.1353966

37. Gernat T, Jagla T, Jones BM, Middendorf M, Robinson GE. Automated monitoring of honey bees with barcodes and artificial intelligence reveals two distinct social networks from a single affiliative behavior. Sci Rep. 2023;13: 1541. doi:10.1038/s41598-022-26825-4

38. Takahashi DY, Baccalá LA, Sameshima K. Information theoretic interpretation of frequency domain connectivity measures. Biol Cybern. 2010;103: 463–469. doi:10.1007/s00422-010-0410-x

39. Sameshima K, Baccalá LA. Methods in brain connectivity inference through multivariate time series analysis. CRC Press; 2014.

40. Hamilton AR, Traniello IM, Ray AM, Caldwell AS, Wickline SA, Robinson GE. Division of labor in honey bees is associated with transcriptional regulatory plasticity in the brain. J Exp Biol. 2019;222: jeb200196. doi:10.1242/jeb.200196

41. Shpigler HY, Saul MC, Murdoch EE, Cash-Ahmed AC, Seward CH, Sloofman L, et al. Behavioral, transcriptomic and epigenetic responses to social challenge in honey bees. Genes, Brain Behav. 2017;16: 579–591. doi:10.1111/gbb.12379

42. Shpigler HY, Saul MC, Murdoch EE, Corona F, Cash-Ahmed AC, Seward CH, et al. Honey bee neurogenomic responses to affiliative and agonistic social interactions. Genes, Brain Behav. 2019;18: e12509. doi:10.1111/gbb.12509

43. Shpigler HY, Robinson GE. Laboratory assay of brood care for quantitative analyses of individual differences in honey bee (*Apis mellifera*) affiliative behavior. PLoS ONE. 2015;10: e0143183. doi:10.1371/journal.pone.0143183

44. Wilson ML. The biology of the honey bee. Harvard University Press; 1991.

45. Robinson GE, Jr. REP, Strambi C, Strambi A. Hormonal and genetic control of behavioral integration in honey bee colonies. Science. 1989;246: 109–112. doi:10.1126/science.246.4926.109

46. Giray T, Robinson GE. Effects of intracolony variability in behavioral development on plasticity of division of labor in honey bee colonies. Behav Ecol Sociobiol. 1994;35: 13–20. doi:10.1007/bf00167054

47. Huang Z-Y, Robinson GE. Regulation of honey bee division of labor by colony age demography. Behav Ecol Sociobiol. 1996;39: 147–158. doi:10.1007/s002650050276

48. Johnson BR. Honey Bee Biology. Princeton University Press; 2023.

49. Lutz CC, Robinson GE. Activity-dependent gene expression in honey bee mushroom bodies in response to orientation flight. J Exp Biol. 2013;216: 2031–2038. doi:10.1242/jeb.084905

50. Browning SR, Browning BL. Rapid and accurate haplotype phasing and missing-data inference for whole-genome association studies by use of localized haplotype clustering. Am J Hum Genet. 2007;81: 1084–1097. doi:10.1086/521987

51. Browning BL, Zhou Y, Browning SR. A one-penny imputed genome from next-generation reference panels. Am J Hum Genet. 2018;103: 338–348. doi:10.1016/j.ajhg.2018.07.015

52. Zheng X, Levine D, Shen J, Gogarten SM, Laurie C, Weir BS. A high-performance computing toolset for relatedness and principal component analysis of SNP data. Bioinformatics. 2012;28: 3326–3328. doi:10.1093/bioinformatics/bts606

53. Gogarten SM, Sofer T, Chen H, Yu C, Brody JA, Thornton TA, et al. Genetic association testing using the GENESIS R/Bioconductor package. Bioinformatics. 2019;35: 5346–5348. doi:10.1093/bioinformatics/btz567

54. Avalos A, Fang M, Pan H, Lluch AR, Lipka AE, Zhao SD, et al. Genomic regions influencing aggressive behavior in honey bees are defined by colony allele frequencies. Proc Natl Acad Sci. 2020;117: 17135– 17141. doi:10.1073/pnas.1922927117

55. Wallberg A, Bunikis I, Pettersson OV, Mosbech M-B, Childers AK, Evans JD, et al. A hybrid de novo genome assembly of the honeybee, Apis mellifera, with chromosome-length scaffolds. BMC Genom. 2019;20: 275. doi:10.1186/s12864-019-5642-0

56. Dobin A, Davis CA, Schlesinger F, Drenkow J, Zaleski C, Jha S, et al. STAR: ultrafast universal RNA-seq aligner. Bioinformatics. 2013;29: 15–21. doi:10.1093/bioinformatics/bts635

57. Liao Y, Smyth GK, Shi W. featureCounts: an efficient general purpose program for assigning sequence reads to genomic features. Bioinformatics. 2014;30: 923–930. doi:10.1093/bioinformatics/btt656

58. Traniello IM, Bukhari SA, Kevill J, Ahmed AC, Hamilton AR, Naeger NL, et al. Meta-analysis of honey bee neurogenomic response links Deformed wing virus type A to precocious behavioral maturation. Sci Rep. 2020;10: 3101. doi:10.1038/s41598-020-59808-4

59. Eden E, Navon R, Steinfeld I, Lipson D, Yakhini Z. GOrilla: a tool for discovery and visualization of enriched GO terms in ranked gene lists. BMC Bioinform. 2009;10: 48. doi:10.1186/1471-2105-10-48

60. Reijnders MJMF, Waterhouse RM. Summary visualizations of Gene Ontology terms with GO-Figure! Front Bioinform. 2021;1: 638255. doi:10.3389/fbinf.2021.638255

61. Elsik CG, Worley KC, Bennett AK, Beye M, Camara F, Childers CP, et al. Finding the missing honey bee genes: lessons learned from a genome upgrade. BMC Genom. 2014;15: 86. doi:10.1186/1471-2164-15-86

62. Gachomba MJM, Esteve-Agraz J, Caref K, Maroto AS, Bortolozzo-Gleich MH, Laplagne DA, et al. Multimodal cues displayed by submissive rats promote prosocial choices by dominants. Curr Biol. 2022;32: 3288–3301.e8. doi:10.1016/j.cub.2022.06.026

63. McNeill MS, Kapheim KM, Brockmann A, McGill TAW, Robinson GE. Brain regions and molecular pathways responding to food reward type and value in honey bees. Genes, Brain Behav. 2016;15: 305–317. doi:10.1111/gbb.12275

64. Naeger NL, Robinson GE. Transcriptomic analysis of instinctive and learned reward-related behaviors in honey bees. J Exp Biol. 2016;219: 3554–3561. doi:10.1242/jeb.144311

65. Wheeler MM, Ament SA, Rodriguez-Zas SL, Southey B, Robinson GE. Diet and endocrine effects on behavioral maturation-related gene expression in the pars intercerebralis of the honey bee brain. J Exp Biol. 2015;218: 4005–4014. doi:10.1242/jeb.119420

66. Lee E-J, Choi SY, Kim E. NMDA receptor dysfunction in autism spectrum disorders. Curr Opin Pharmacol. 2015;20: 8–13. doi:10.1016/j.coph.2014.10.007

67. Jan LY, Jan YN. L-glutamate as an excitatory transmitter at the *Drosophila* larval neuromuscular junction. J Physiol. 1976;262: 215–236. doi:10.1113/jphysiol.1976.sp011593

68. Si A, Helliwell P, Maleszka R. Effects of NMDA receptor antagonists on olfactory learning and memory in the honeybee (Apis mellifera). Pharmacol Biochem Behav. 2004;77: 191–197. doi:10.1016/j.pbb.2003.09.023

69. Varoqueaux F, Jamain S, Brose N. Neuroligin 2 is exclusively localized to inhibitory synapses. Eur J Cell Biol. 2004;83: 449–456. doi:10.1078/0171-9335-00410

70. Ali H, Marth L, Krueger-Burg D. Neuroligin-2 as a central organizer of inhibitory synapses in health and disease. Sci Signal. 2020;13: eabd8379. doi:10.1126/scisignal.abd8379

71. Parente DJ, Garriga C, Baskin B, Douglas G, Cho MT, Araujo GC, et al. Neuroligin 2 nonsense variant associated with anxiety, autism, intellectual disability, hyperphagia, and obesity. Am J Méd Genet Part A. 2017;173: 213–216. doi:10.1002/ajmg.a.37977

72. Corthals K, Heukamp AS, Kossen R, Großhennig I, Hahn N, Gras H, et al. Neuroligins Nlg2 and Nlg4 affect social behavior in *Drosophila melanogaster*. Front Psychiatry. 2017;8: 113. doi:10.3389/fpsyt.2017.00113

73. Paffhausen BH, Fuchs I, Duer A, Hillmer I, Dimitriou IM, Menzel R. Neural correlates of social behavior in mushroom body extrinsic neurons of the honeybee *Apis mellifera*. Front Behav Neurosci. 2020;14: 62. doi:10.3389/fnbeh.2020.00062

74. Fischer EK, O’Connell LA, Levine JD, Kronauer DJC, Dickinson MH. Modification of feeding circuits in the evolution of social behavior. J Exp Biol. 2017;220: 92–102. doi:10.1242/jeb.143859

75. Fitzpatrick P, Romero V, Amaral JL, Duncan A, Barnard H, Richardson MJ, et al. Evaluating the importance of social motor synchronization and motor skill for understanding autism. Autism Res. 2017;10: 1687–1699. doi:10.1002/aur.1808

76. McNaughton KA, Redcay E. Interpersonal synchrony in autism. Curr Psychiatry Rep. 2020;22: 12. doi:10.1007/s11920-020-1135-8

77. Dridi S, Akçay E. Learning to cooperate: The evolution of social rewards in repeated interactions. Am Nat. 2018;191: 58–73. doi:10.1086/694822

78. Walter H, Abler B, Ciaramidaro A, Erk S. Motivating forces of human actions: Neuroimaging reward and social interaction. Brain Res Bull. 2005;67: 368–381. doi:10.1016/j.brainresbull.2005.06.016

79. Huang J, Zhang Z, Feng W, Zhao Y, Aldanondo A, Sanchez MG de B, et al. Food wanting is mediated by transient activation of dopaminergic signaling in the honey bee brain. Science. 2022;376: 508–512. doi:10.1126/science.abn9920

80. Liang ZS, Nguyen T, Mattila HR, Rodriguez-Zas SL, Seeley TD, Robinson GE. Molecular determinants of scouting behavior in honey bees. Science. 2012;335: 1225–1228. doi:10.1126/science.1213962

81. Lehman HK, Schulz DJ, Barron AB, Wraight L, Hardison C, Whitney S, et al. Division of labor in the honey bee (*Apis mellifera*): the role of tyramine β-hydroxylase. J Exp Biol. 2006;209: 2774–2784. doi:10.1242/jeb.02296

82. Tedjakumala SR, Rouquette J, Boizeau M-L, Mesce KA, Hotier L, Massou I, et al. A tyrosine- hydroxylase characterization of dopaminergic neurons in the honey bee brain. Front Syst Neurosci. 2017;11: 47. doi:10.3389/fnsys.2017.00047

83. Li Q, Wang M, Zhang P, Liu Y, Guo Q, Zhu Y, et al. A single-cell transcriptomic atlas tracking the neural basis of division of labour in an ant superorganism. Nat Ecol Evol. 2022;6: 1191–1204. doi:10.1038/s41559-022-01784-1

84. Sheng L, Shields EJ, Gospocic J, Glastad KM, Ratchasanmuang P, Berger SL, et al. Social reprogramming in ants induces longevity-associated glia remodeling. Sci Adv. 2020;6: eaba9869. doi:10.1126/sciadv.aba9869

